# Remdesivir is a delayed translocation inhibitor of SARS CoV-2 replication in vitro

**DOI:** 10.1101/2020.12.14.422718

**Authors:** Jack P. K. Bravo, Tyler L. Dangerfield, David W. Taylor, Kenneth A. Johnson

## Abstract

Remdesivir is a nucleoside analog approved by the FDA for treatment of COVID-19. Here, we present a 3.9-Å-resolution cryoEM reconstruction of a remdesivir-stalled RNA-dependent RNA polymerase complex, revealing full incorporation of three copies of remdesivir monophosphate (RMP) and a partially incorporated fourth RMP in the active site. The structure reveals that RMP blocks RNA translocation after incorporation of three bases following RMP, resulting in delayed chain termination, which can guide the rational design of improved antiviral drugs.

## Introduction

Although vaccines appear to be on track to mitigate the coronavirus disease 2019 (COVID-19) pandemic, there is still a need for direct acting antiviral drugs to treat severe acute respiratory syndrome coronavirus 2 (SARS-CoV-2) infections when immunization fails and for future coronavirus outbreaks.

Structures of the core SARS-CoV-2 RNA-dependent RNA polymerase (RdRp) complex (consisting of the enzyme nsp12 and the accessory subunits nsp7 and nsp8) have revealed the structural basis for RNA replication (Chen et al., 2020; Hillen et al., 2020; Wang et al., 2020b; Yan et al., 2020a; Yan et al., 2020b; Yin et al., 2020). However, the mechanism of action of remdesivir, an adenosine analog that was recently approved by the FDA for COVID-19 treatment, remains to be established (Beigel et al., 2020; Wang et al., 2020a). Kinetic studies have shown the remdesivir triphosphate (RTP) is incorporated rapidly with efficiency two-fold greater than ATP and is rapidly extended within 1 s, but the polymerase stalls after adding two or three nucleotides on top of the newly incorporated remdesivir monophosphate (RMP) (Dangerfield et al., 2020; Gordon et al., 2020). Here, we provide the structure of a stalled SARS-CoV-2 polymerase-RNA complex stalled after incorporation of remedesivir, revealing the underlying mechanistic basis for inhibition of replication.

## Results

### Kinetics of remdesivir incorporation and stalling by RdRp

We first examined the kinetics of remdesivir incorporation and subsequent stalling of continued RNA replication by incubating the RdRp with a FAM-labeled primer:template RNA duplex (Figure 1A) and then initiating the polymerization by addition of uracil triphosphate (UTP) and RTP (in the presence of 5 mM Mg^2+^). After quenching the reaction with EDTA, we resolved and quantified the reaction products at different time points using capillary electrophoresis (CE) by monitoring fluorescence. While we have previously demonstrated rapid incorporation of 2 UTP and 3 RTP molecules within 1 s (Dangerfield et al., 2020), here we found that over the time course of 20 s, there is a significantly slower incorporation of a fourth RTP (a rate of 0.13 s^−1^, at the concentrations listed in Figure 1B). Note that we did not observe further addition of UTP on top of the 4 RMPs observed under these conditions, indicating that RdRp complex has stalled (Figure 1C). CE reveals a mixture of 25- and 26-nt products corresponding to the extended primer with a mixture of 3 (10%) and 4 RMP (90%) molecules incorporated, respectively (Figure 1C). Under the same conditions in the presence of UTP and ATP, we see the extension to form a 27 nt product, indicating that premature stalling does not occur with ATP (Figure 1D).

**Figure 1.**
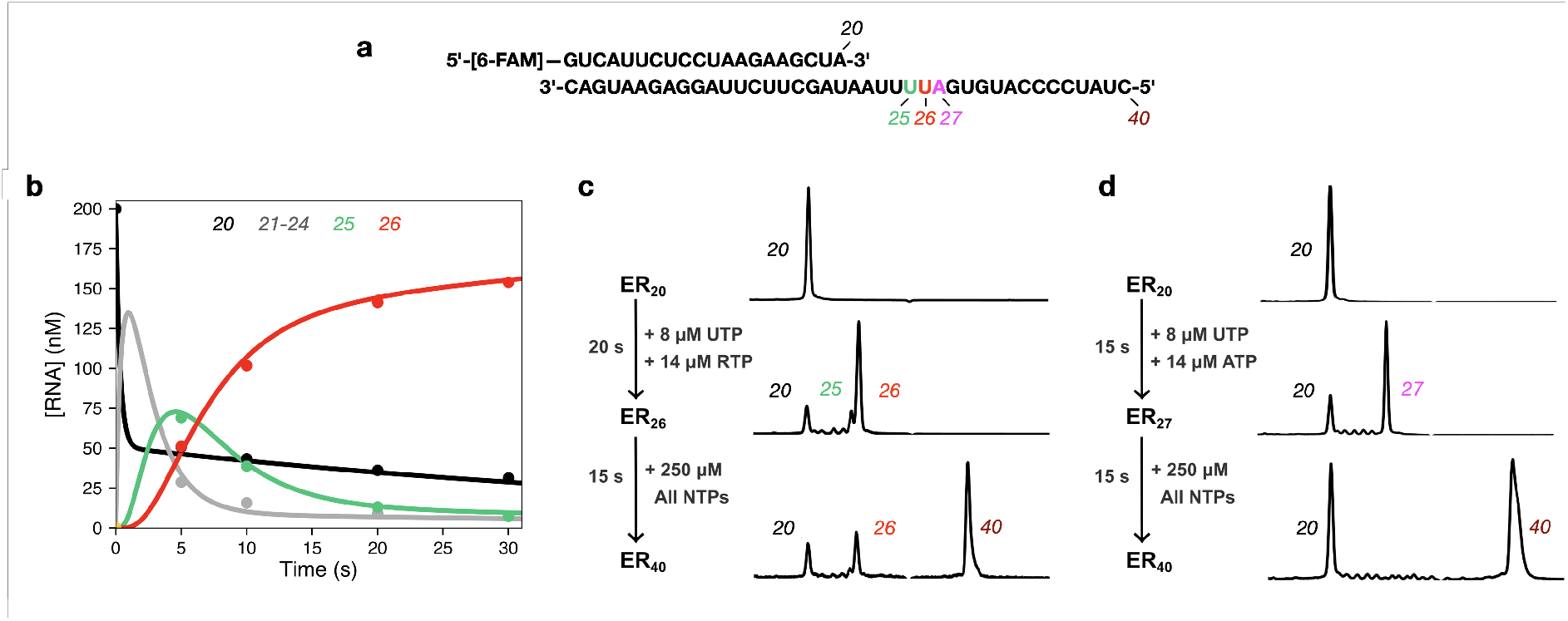
RdRp stalls after incorporation of 4 remdesivirs. (A) The RNA substrate used in experiments consists of a 20 nt, 5’-[6-FAM] labeled primer annealed to a 40 nt template. The sequence is from the 3’ end of the SARS-CoV-2 genome, preceding the poly-A tail. (B) Time course of UTP + RTP incorporation by the RdRp complex. A solution of 3.33 μM NSP12/7/8, 4.7 μM NSP8 and 200 nM FAM-20/40 RNA was mixed with 14 μM RTP and 8 μM UTP to start the reaction. Time points were quenched by addition of EDTA and samples were analyzed by capillary electrophoresis. Lengths of various RNA products are given at the top of the panel. Solid lines through the data points are from the fit by simulation in KinTek Explorer. (C) UTP + RTP product formation and extension. The chromatogram for the zero-time point is given at the top (ER_20_). Initially RTP and UTP were added for 20 seconds (ER_26_) at the concentrations given in the figure. Then 250 μM of all NTPs were added for 15 seconds to allow extension to the full-length product (ER_40_). On the timescale of the experiment, only a fraction of ER_26_ gets converted to full length product (ER_40_). (D) UTP + ATP product formation and extension. The experiment was performed as in (C), but with ATP instead of RTP in the first mixing step, allowing extension to the ER27 product. After addition of all nucleotides for 15 seconds in the second mixing step, the ER27 complex is completely converted to full length ER_40_ product.

Following the addition of 250 μM of all four nucleotides to the 27-nt product formed with ATP, we observed complete extension to the full-length 40-nt product within 15s. In contrast, when we add 250 μM of all four nucleotides to the RMP-stalled RNA, only 70% of the 26-nt (RTP) primer was fully extended to yield a 40-nt product within 15 s. The apparent extension rate of 0.08 s^−1^ represents a 4000-fold inhibition of normal incorporation. While our results are in agreement with previous studies that have shown that RMP incorporation results in stalling after being extended by three nucleotides, our data collected with a highly active replicating enzyme show that the inhibition is not absolute and can be overcome by incubation at high nucleotide concentrations.

### A physical barrier between remdesivir and Rdrp blocks translocation

We used conditions that lead to stalling in our kinetic analysis above to prepare samples for cryo-EM, allowing us to determine the structure of *a bona fide* elongation intermediate at a global resolution of 3.9 Å (Figure 2). We observe well-resolved primer:template duplex density, allowing *de novo* modelling of a remdesivir-stalled primer elongation intermediate. Within the active site of the RdRp complex, remdesivir monophosphate (RMP) occupies positions −3, −2, and −1 (i.e., 3, 2, and 1 nucleotides downstream from the active site, respectively). We also observed additional density in the polymerase active site (position +1) position. This density corresponds to the nucleobase of a fourth remdesivir, consistent with our kinetic analysis (Figure 1C). This suggests that while the fourth RMP can participate in Watson-Crick basepairing with the template UMP and stacking interactions with neighboring RMP, the sugarphosphate backbone may be somewhat disordered. Surprisingly, the fourth RMP was retained at the polymerase active site and did not translocate to allow the binding of the next nucleotide (UTP in our sequence). Thus, our data suggest that the polymerase stalls because the translocation step is blocked. We modelled a hypothetical post-translocated state, where the first RMP is moved to the −4 position, which revealed a steric clash between the C1 cyano group of RMP(−4) and the side-chain of Ser861 of the nsp12 (Figure 2E-H). This residue provides a physical barrier to slow duplex translocation and may lead to perturbations near or in the active site that slow the rate of chain elongation. Our model provides a structural basis for stalling of RdRp after incorporation of the fourth RMP. This is in agreement with previous studies that have demonstrated that mutations of nsp12(Ser861) reduce the inhibitory effect of remdesivir (Gordon et al., 2020; Tchesnokov et al., 2020). Our structure also shows that the cyano group of each of the four RMP molecules is readily accommodated at the active site, with no apparent steric interactions with the enzyme. The proposed steric interaction would only occur as the RNA duplex attempts to translocate to allow binding and incorporation of UTP.

**Figure 2.**
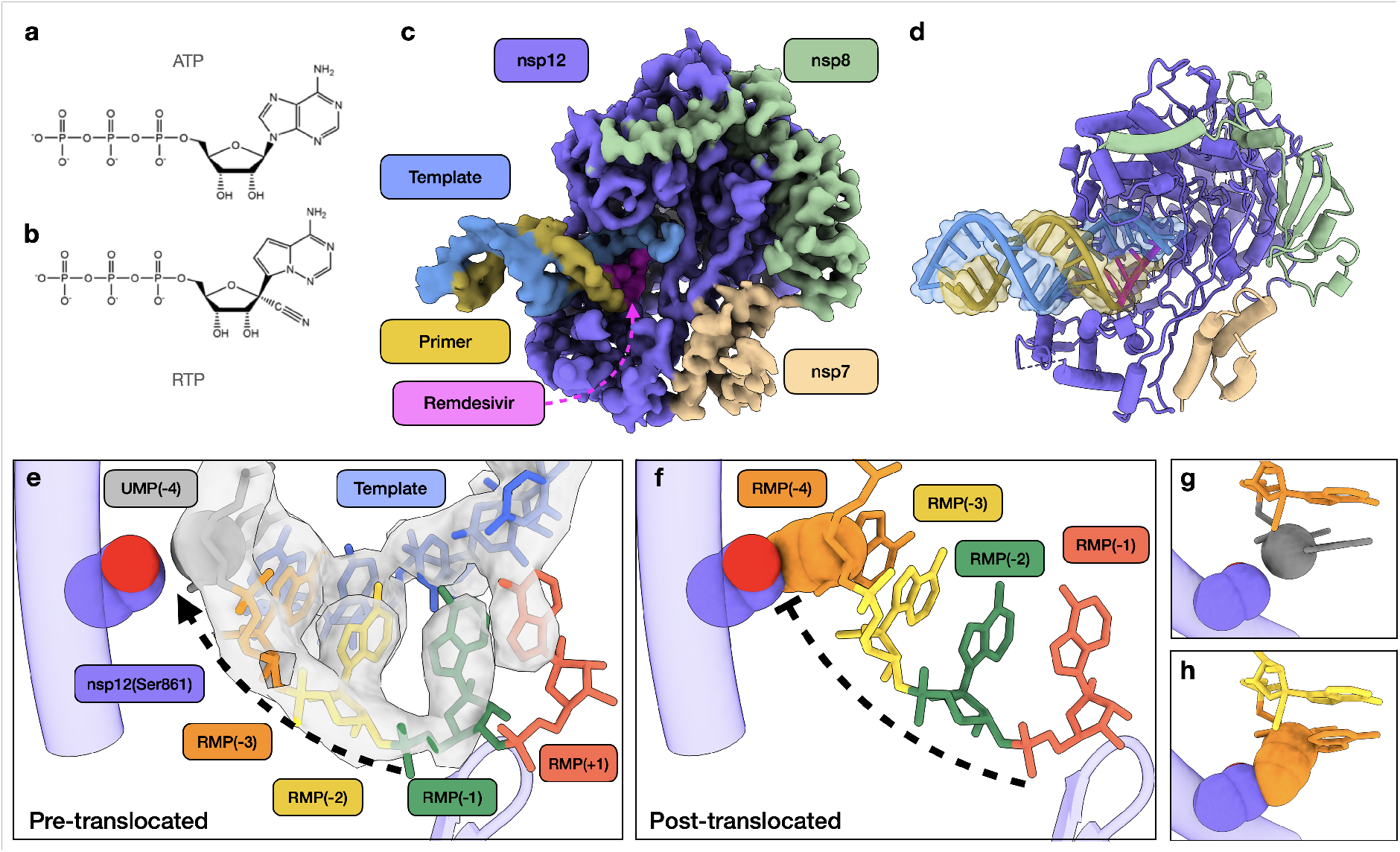
Structural basis of SARS-CoV-2 RdRp complex stalling by remdesivir. (A) Chemical structures of adenosine triphosphate. (B) Chemiccal structure of remdesivir triphosphate. (C) Cryo-EM reconstruction of remdesivir-stalled RdRp complex. (D) Corresponding atomic model of remdesivir-stalled RdRp complex. (E) Close-up view of polymerase active site. Four copies of remdesivir monophosphate (RMP) occupy positions +1 (active site), −1, −2, −3. Uracil monophosphate (UMP) is in position −4 (grey). C1 carbon of UMP and side-chain of nsp12(Ser861) are shown as spheres. CryoEM density corresponding to template RNA strand (blue) and primer strand positions +1 through −5 is shown (transparent grey). Active site loop is also shown. (F) Hypothetical model of post-translocated state, whereby the first copy of RMP is translocated into the −4 polymerase position. RMP(−4) cyano group and nsp12(Ser861) are shown as spheres, and show a steric clash. (G) Spatial relationship between nucleotide at −4 position and side-chain of nsp12(Ser861) in pre-translocated state. (H) Spatial relationship between nucleotide at −4 position and side-chain of nsp12(Ser861) illustrating the clash between the cyano group of RMP that prevents translocation. See Table S1 and Figure S1 for cryo-EM data collection, model validation, and analysis.

## Discussion

Combined with prior our kinetic analysis, our structure suggests why the SARS CoV-2 RdRp can rapidly incorporate three molecules of RMP then more slowly incorporate a fourth RMP, which remains in the active site. This demonstrates that the RNA fails to be translocated to open the active site to allow binding of the next nucleotide. Translocation is inhibited by the steric interaction between Ser861 and the cyano group of the first RMP (seen in the structure at the −4 position). Interpretation of the structural data together with the kinetic data define the mechanism of inhibition by remdesivir as a delayed chain extension inhibitor. We propose that the steric block between the cyano group of RMP at −4 slows translocation to inhibit continued polymerization. This inhibition is not absolute, in that subsequent addition of a high concentration of the four nucleotides supports extension of 70% of the RNA within 20 s.

While there have been multiple structures of SARS-CoV-2 with a single copy of RMP incorporated determined (Kokic, 2020; Wang et al., 2020b; Yin et al., 2020), our structure reveals the first insights into a *bona fide* stalled elongation complex of an actively polymerizing enzyme. Initial structural studies with a single copy of RMP in positions −1 (Wang et al., 2020b) and +1 (Yin et al., 2020) demonstrate that RMP is able to be incorporated into the nascent transcript through Watson-Crick base pairing with template strand uracil bases. However, together with previous studies (Gordon et al., 2020; Tchesnokov et al., 2020), our structure and kinetic data show that incorporation of RMP at neither of these positions results in RdRp inhibition. Another recent study (Kokic, 2020) determined multiple structures of RdRp in complex with synthetic substrates in which RMP had been incorporated at specific positions. While both that study and ours have identified a clash between RMP(−4) and nsp12(Ser861) as the translocation barrier inhibiting elongation and thus inducing RdRp stalling, our additional analysis reveals that this state represents a kinetically trapped intermediate.

These studies were performed in the absence of the proofreading exonuclease (nsp10/14), and as such, we cannot fully define the fate of remdesivir and subsequent elongation products that will be observed with the complete holoenzyme. Exonuclease selectivity is determined at the polymerization site by stalling of the polymerase to slow extension of a mismatched base pair, which then serves as a trigger for exonuclease removal by altering the kinetic partitioning between extension versus excision (Donlin et al., 1991). Stalling of the polymerase gives time for the transfer of the primer strand from the polymerase to the exonuclease site. It is tempting to hypothesize that RMP may be resistant to excision, not because of inherent properties of RMP, but rather because subsequent extension after the first RMP incorporation (Dangerfield et al., 2020) is faster than expected rates of primer strand transfer and excision at the exonuclease site (Donlin et al., 1991; Johnson and Johnson, 2001). Thus, kinetic partitioning favors extension rather than excision of the first RMP.

The steric clash between serine and cyano group of RMP does not completely block polymerization, so over time the RNA is extended (at a net rate of 0.08 s^−1^) to bury the incorporated RMP in the absence of the proofreading exonuclease. After the addition of three nucleotides on top of the first RMP, stalling of the polymerase is expected to lead to exonuclease removal of the 3’-terminal base. According to this kinetic model, the excised base would then be rapidly replaced, and could lead to a steady-state balance with repeated cycles of excision and replacement of the 3’-terminal base pair, effectively blocking further extension while protecting RMP from excision. Remdesivir is five times more effective in viral mutants with reduced exonuclease activity (Agostini et al., 2018), so it is clear that the exonuclease reduces the effectiveness of remdesivir. In the absence of the exonuclease, remdesivir could become buried in the genome which could cause lethal mutagenesis (Shannon et al., 2020). Further studies are required to establish the kinetic parameters and structural features that govern exonuclease proofreading relative to RMP-induced polymerase stalling. Nonetheless, our current data provide a structural basis for RMP-induced stalling, which may guide the development of more effective nucleoside analogs.

## Methods

### Material preparation: protein, RNA substrates, and other materials

The SARS CoV-2 RdRp (nsp12/7/8) was expressed and purified without tags to provide a highly active enzyme for transient kinetic and structural studies using cryo-electron microscopy (cryo-EM). Tag-free NSP12/7/8 complex and NSP8 were expressed in *E. coli* and purified as described previously (Dangerfield et al., 2020). RNA samples were prepared in DEPC treated water (Ambion). RNA substrates were purchased from Integrated DNA technologies with RNase free HPLC purification and resuspended in Annealing Buffer (10 mM Tris-HCl pH 7, 50 mM NaCl, 0.1 mM EDTA). Concentration of each oligo was determined by absorbance at 260 nm using the extinction coefficients 222,360 M^−1^ cm^−1^ and 403,100 M^−1^ cm^−1^ for the FAM-20 nt primer and the 40 nt template, respectively. The double stranded RNA substrate was prepared by mixing each oligo at a 1:1 molar ratio, heating to 75°C for 3 minutes, then cooling slowly to room temperature over approximately 2 hours. Oligonucleotides were stored at −20°C. UTP was purchased from New England Biolabs. Remdesivir triphosphate (GS-443902) was a kindly provided by Gilead Sciences and concentration was determined by absorbance at 245 nm using the extinction coefficient 24,100 M^−1^ cm^−1^ (Dangerfield et al., 2020). Remdesivir triphosphate was stored at −20°C.

### Extension reactions and analysis by capillary electrophoresis

All reactions were conducted in Reaction Buffer (40 mM Tris-HCl pH 7, 50 mM NaCl, 5 mM MgCl2, 1 mM DTT) (Dangerfield et al., 2020). Concentrations of enzyme, RNA and nucleotides are given in the legend for Figure 1. (concentrations given are the final concentrations after mixing). Reactions were performed at room temperature (~25°C) in Reaction Buffer after allowing the RdRp complex and RNA to equilibrate for approximately 30 minutes. Samples were quenched by adding 0.6 M EDTA to a final concentration of 0.4 M. One μL of each sample was diluted into 10 μL of HiDi formamide (ThermoFisher) containing 1 nM Cy3 internal standard DNA oligo (5’-[Cy3]-CCGTGAGTTGGTTGGACGGCTGCGAGGC-3’, purchased from Integrated DNA Technologies) in a 96 well plate. Samples were analyzed on an Applied Biosystems 3130xl Genetic Analyzer instrument equipped with a 36 cm capillary array (ThermoFisher) and nanoPOP-6 polymer (Molecular Cloning Laboratories). The oven temperature was set to 65°C. Before injecting samples, a pre-run electrophoresis step was performed at 15 kV for 3 minutes. Samples were injected at 3.6 kV for 12 seconds, then the voltage was ramped up to 15 kV over 40, 15 second steps. Fluorescence was monitored for 800 seconds using the G5 dye set and all peak intensities were within the linear range of the instrument. Peaks were integrated with GeneMapper 5 software (ThermoFisher).

### Cryo-EM sample preparation, data collection and processing

Samples were prepared in Reaction Buffer. A solution of 3.33 μM NSP12/7/8, 4.67 μM NSP8, 3.33 μM FAM-20/40 RNA was mixed with 14 μM Remdesivir Triphosphate (RTP) and 7.5 μM UTP to start the reaction at room temperature (~25°C) (concentrations of reaction components are given after mixing). The reaction was allowed to proceed for 10-20 seconds before application to glow discharged holey carbon grids (C-flat 4/2, Protochips Inc.), blotted for 0.5 s with a blot force of 4 and rapidly plunged into liquid ethane using an FEI Vitrobot MarkIV.

Data was collected on an FEI Titan Krios cryo-electron microscope equipped with a K3 Summit direct electron detector (Gatan, Pleasanton, CA). Images were recorded with SerialEM (Mastronarde, 2005), with a pixel size of 1.1Å over a defocus range of −1.5 to −2.5 μm. 2446 movies were recorded at 13.3 electrons/pixel/second for 6 seconds (80 frames) to give a total dose of 80 electrons/pixel. CTF correction, motion correction and particle picking were performed in real-time using WARP, resulting in 2,340,544 particles, which were uploaded to cryoSPARC v2 (Punjani et al., 2017).

Particles were subjected to multiple rounds of 3D classification in cryoSPARC, and a final set of 116,748 particles was refined to a global resolution of 3.89 Å based on the 0.143 FSC criterion. Map sharpening was performed using DeepEMhancer (Sanchez-Garcia et al, 2020) as implemented in COSMIC^2^ (Cianfrocco et al 2017). For modelling, the published structure of nsp7-8-12 (PDB: 7bv2 (Yin et al., 2020)) was used as a starting model. Non-proteinaceous molecules (i.e. RNA, ligands) were removed, and the template:primer duplex was built de novo in Coot (Emsley and Cowtan, 2004). Restraints for RMP were generated using eLBOW (Moriarty et al., 2009)), and structures were subjected to real-space refinement using Phenix (which resulted in a map-to-model FSC of 4.0Å at the 0.5 threshold) (Afonine et al., 2018). Figures were prepared using ChimeraX (Goddard et al., 2018).

## Author Contributions

J.P.K.B. performed cryo-EM, structure determination, and modeling. T.L.D. purified the enzyme, performed kinetic studies and prepared samples for cryo-EM. J.P.K.B., T.L.D., D.W.T., and K.A.J. analyzed and interpreted the data and wrote the manuscript. D.W.T. and K.A.J. supervised and secured funding for the studies.

## Data Availability

The cryo-EM structure and associated atomic coordinates are available at the Electron Microscopy Data Bank and Protein Data Bank with accession codes EMD-23109 and PDB ID 7L1F, respectively.

## Acknowledgements

We thank Gilead (assisted by Brian Schultz and Joy Feng) for providing the Remdesivir triphosphate used in this study. This work was supported in part by Welch Foundation grants F-1604 (to K.A.J.) and F-1938 (to D.W.T.), National Institute of Allergy and Infectious Diseases (NIAID) of the National Institutes of Health (NIH) R01AI110577 (to K.A.J) and National Institute of General Medical Sciences (NIGMS) of the National Institutes of Health (NIH) R01114223 (to K.A.J.) and R35GM138348 (to D.W.T.), Army Research Office Grant W911NF-15-1-0120 (to D.W.T.), and a Robert J. Kleberg, Jr. and Helen C. Kleberg Foundation Medical Research Award (to D.W.T.). D.W.T is a CPRIT Scholar supported by the Cancer Prevention and Research Institute of Texas (RR160088) and an Army Young Investigator supported by the Army Research Office (W911NF-19-1-0021).

---

**Table S1.**
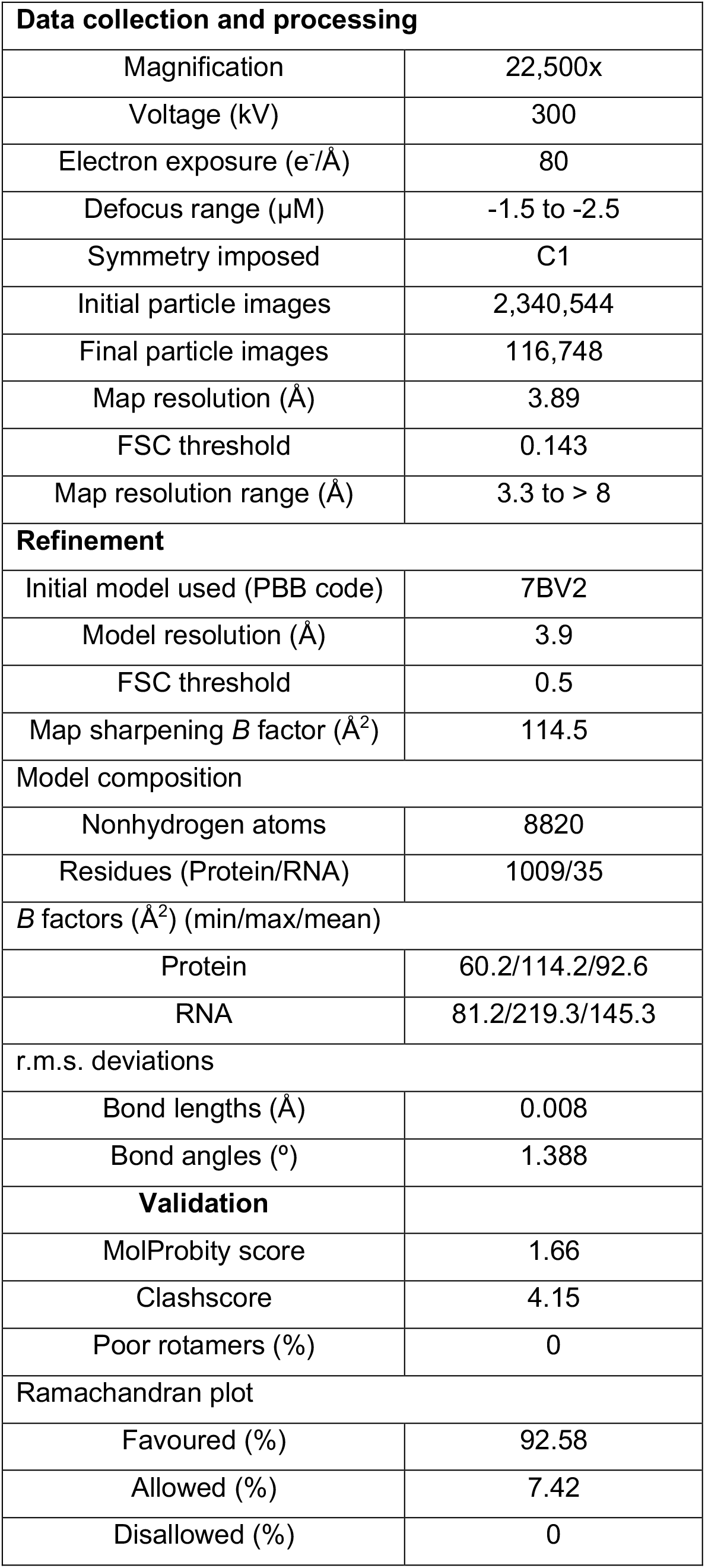
Cryo-EM data collection and processing parameters. Related to Figure 2.

| Data collection and processing |  |
| --- | --- |
| Magnification | 22,500x |
| Voltage (kV) | 300 |
| Electron exposure (e<sup>-</sup>/Å) | 80 |
| Defocus range (μM) | -1.5 to -2.5 |
| Symmetry imposed | C1 |
| Initial particle images | 2,340,544 |
| Final particle images | 116,748 |
| Map resolution (Å) | 3.89 |
| FSC threshold | 0.143 |
| Map resolution range (Å) | 3.3 to > 8 |
| Refinement |  |
| Initial model used (PBB code) | 7BV2 |
| Model resolution (Å) | 3.9 |
| FSC threshold | 0.5 |
| Map sharpening *B* factor (Å<sup>2</sup>) | 114.5 |
| Model composition |  |
| Nonhydrogen atoms | 8820 |
| Residues (Protein/RNA) | 1009/35 |
| *B* factors (Å<sup>2</sup>) (min/max/mean) |  |
| Protein | 60.2/114.2/92.6 |
| RNA | 81.2/219.3/145.3 |
| r.m.s. deviations |  |
| Bond lengths (Å) | 0.008 |
| Bond angles (°) | 1.388 |
| Validation |  |
| MolProbity score | 1.66 |
| Clashscore | 4.15 |
| Poor rotamers (%) | 0 |
| Ramachandran plot |  |
| Favoured (%) | 92.58 |
| Allowed (%) | 7.42 |
| Disallowed (%) | 0 |

**Figure S1.**
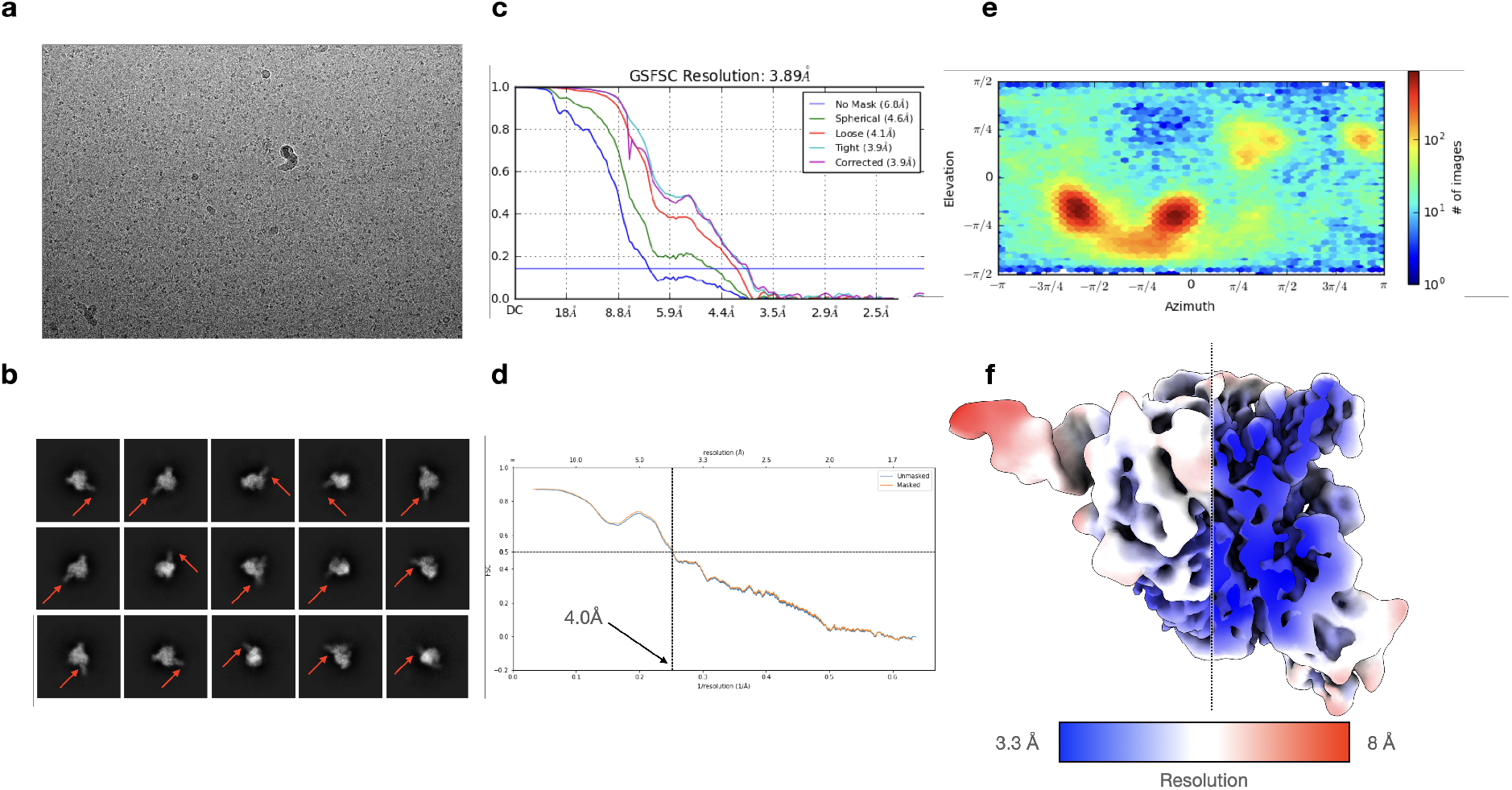
Cryo-EM analysis of stalled RdRp complex. Related to Figure 2. (A) Representative cryo-EM micrograph of vitrified nsp7-8-12:RNA:remdesivir. (B) Representative 2D class averages, with extended template:primer duplex denoted by red arrow. (C) Gold-standard FSC of final reconstruction, indicating a global resolution of 3.9Å at the 0.143 threshold. (D) Map-to-model FSC, showing a resolution of 4.0Å at the 0.5 threshold. (E) Euler angle distribution of the final reconstruction. (F) Final reconstruction filtered and colored according to local resolution. The core of the complex (including the RMP-containing active site) is at a local resolution of ~3.3 Å.

